# Spatial Bias in Lesion Network Mapping Is Connectome-Independent

**DOI:** 10.64898/2026.03.17.712378

**Authors:** Max Wawrzyniak, Tim Ritter, Julian Klingbeil, Gordian Prasse, Dorothee Saur, Anika Stockert

## Abstract

Lesion network mapping (LNM) is increasingly used to link focal brain lesions to distributed functional networks. Recent work has raised concerns that LNM results may be spatially biased by dominant features of the normative connectome.

If this were the case, three testable predictions would follow: (i) a consistent spatial pattern of false positives across LNM studies, (ii) that this pattern can be consistently explained by intrinsic connectome organization, and (iii) that symptom-associated LNM findings preferentially occur in regions with high spatial bias.

We tested these predictions across three independent LNM datasets (n = 49/101/200), evaluating each prediction in all cohorts. Spatial bias maps derived from 4,000,000 random permutations under the null hypothesis showed minimal correspondence across cohorts (R^2^ = 0.4–0.8%), indicating strong cohort specificity. Moreover, dominant connectome features—captured by the first 10 principal components of connectivity profiles from 1,000 atlas regions—did not systematically explain these bias maps. Finally, symptom-associated results showed no enrichment in high-bias regions.

Together, these findings provide strong evidence that spatial bias in LNM is not driven by dominant connectome features. With appropriate inferential statistics and rigorous study design, LNM remains a valid approach for mapping symptom-related brain networks.

## Introduction

Complex behavior emerges from distributed, functionally connected brain networks.^1^ This view shifted clinical neuroscience from a localist perspective toward circuit-based models of dysfunction, with potential implications for a future era of network-informed therapeutic interventions.^2; 3^ Lesion network mapping (LNM) and related connectome-based network analyses adopt this perspective by modeling that neurological, neuropsychological or psychiatric disorders reflect dysfunction within common brain networks rather than solely focal effects of damage. Normative resting-state functional MRI connectome data of healthy individuals is used to estimate the brain-wide functional connectivity profile of individual, heterogeneous focal brain alterations (e.g. lesions, atrophy, activation peaks, stimulation sites). Group-level analyses then link alteration-derived connectivity profiles to behavioral or clinical phenotypes across individuals.^4–6^ In this way LNM provides a unique non-invasive framework for identifying distributed, functionally connected brain regions that may contribute to a behavior, disorder or symptom expression. It has been widely adopted due to its accessibility–requiring only behavioral characterization, maps or coordinates of focal brain alterations and publicly available connectome data. However, to ensure meaningful results and clinical relevance, careful validation of potential methodological biases is essential, particularly when findings are used to support mechanistic interpretations or to derive testable therapeutic targets.

Group-level analyses in LNM are characterized by considerable methodological heterogeneity and have been the subject of debate.^7; 8^ Recently, van den Heuvel and colleagues raised concerns that LNM may be biased by fundamental spatial properties of the normative connectome that could introduce spatial bias. Their critique is grounded in both empirical and analytical observations. Empirically, a high degree of similarity has been reported across LNM studies. Analytically, it has been argued that, by design, LNM effectively samples only basic properties of the normative connectome (e.g., the mean connectivity pattern or dominant principal components) independent of symptom-specific effects. Consequently, the biological specificity and interpretability of LNM results have been questioned.^9^

Several counterarguments have been put forward, including that the critique of van den Heuvel and colleagues reflects an incomplete implementation of the method lacking specificity testing, that many comparisons rely on spatial similarity of unthresholded effect maps, and that spatial similarity alone does not preclude biologically meaningful differences.^10^ We contend that, in evaluating the methodological validity of LNM and the possibility of spatial bias, it is crucial to distinguish clearly between different levels of statistical analysis. Therefore, Table 1 summarizes commonly used descriptive and inferential approaches without claiming to be exhaustive.

**Table 1.** Descriptive and inferential statistical methods in lesion network mapping. References are exemplary and not comprehensive. ^4^ Boes et al., 2015, ^11^Wawrzyniak et al., 2018,^12^Klingbeil et al., 2020, ^13^Stockert et al., 2023, ^14^Schaper et al., 2023, ^15^Al-Fatly et al., 2024, ^16^Klingbeil et al., 2021, ^17^Rangus et al., 2024, ^18^ Sutterer et al., 2016, ^19^Siddiqi et al., 2024.

|  | Method | Level | Control condition | Ref. |
| --- | --- | --- | --- | --- |
| 1 | Sum or mean of LNMs of interest | Descriptive | None | 4; 15; 18 |
| 2 | Calculating (weighted) difference between LNMs of interest and control LNMs | Descriptive | 1. Synthetic lesions<br>2. Lesions from external source<br>3. lesions from patients in control condition | 18 |
| 3 | Weighting LNMs of interest with behavior | Descriptive | Implicit model-based control | 19 |
| 4 | Testing LNMs of interest against zero | Inferential | None | 15 |
| 5 | Testing LNMs of interest against control LNMs | Inferential | 1. Synthetic lesions<br>2. Lesions from external source<br>3. Lesions from patients in control condition | 4; 11–16 |
| 6 | Testing LNMs of interest for correlation with a behavioral variable | Inferential | Implicit model-based control | 14; 17 |

Van den Heuvel et al. demonstrated that the risk of sampling only basic properties of the connectome applies to descriptive analyses and to inferential approaches that lack appropriate control conditions (Table 1, rows 1–4).^9^ However, more rigorous approaches (Table 1, rows 5–6), which combine inferential statistics with appropriate control conditions, were not sufficiently addressed.^11–17^ This leaves open the question of whether LNM is inherently methodologically flawed or concerns instead primarily apply to inappropriate group-level statistical design.

Here, we address this gap by evaluating multiple LNM datasets with control groups using robust inferential statistics (corresponding to row 5.3 in Table 1). We test whether a systematic spatial bias driven by basic properties of the underlying connectome persists in this case and whether results in real-world LNM data can be explained by such a bias.

## Material and methods

### Conceptual framework

A rigorous study design for LNM involves a consecutive cohort of patients with focal brain lesions, characterized with respect to a predefined clinical symptom or behavior. Patients are stratified into symptom+ and symptom− groups. This design mirrors classical lesion–symptom mapping while extending inference from focal anatomy to distributed functional networks.^20^ For each patient, an individual LNM is computed and group-level differences are assessed by comparing LNMs between symptom+ and symptom− patients.

Two methodological issues must be considered here: First, lesion network maps are spatially autocorrelated; second, they are not statistically independent, as all are derived from the same normative connectome. Together, these properties violate assumptions of classical parametric tests. One approach to address these issues—also employed in the present study—is permutation-based inference within the general linear model. By randomly permuting group labels (symptom +/−), an empirical null distribution is generated that preserves the spatial structure of the lesion network data without relying on parametric assumptions. This framework defines the significance threshold for network-symptom associations and allows control of the family-wise error rate across voxels, providing robust inference on group differences of LNM patterns.^21^

According to the hypothesis advocated by van den Heuvel et al., LNM may primarily reflect fundamental spatial properties of the connectome rather than symptom-specific effects. If this assumption holds, we would expect three converging observations to occur simultaneously: (i) a consistent spatial pattern of false positives (i.e., the top 5% of the permutative null distribution) across studies, (ii) that this pattern is persistently explainable by intrinsic connectome organization, and (iii) that symptom-associated findings preferentially fall in regions with high spatial bias.

To test this, the spatial distribution of false positives (spatial bias) under the null hypothesis was empirically sampled using a large number of random permutations based on data from real LNM studies. We then assessed similarity across cohorts and examined the extent to which the observed spatial bias could be explained by dominant features of the underlying connectome. In addition, we displayed the localization of symptom-associated findings within the distribution of false positives to identify potential associations between real-world LNM results and spatial bias.

### Patients and behavioral testing

The analyses were based on three LNM studies on anosognosia for hemiplegia^12^, thalamic aphasia^13^ and post-stroke epilepsy (unpublished). Table 2 gives an overview on demographic information, behavioral testing and LNM procedure. Lesion distributions can be found in SI Figure 1. All details of the LNM procedures can be found in the original publications.^12; 13^ For the post-stroke epilepsy cohort, processing and normative cohort details are identical to another prior study.^16^

**Table 2.** Demographic, behavioral and methodological information. ^22^Placencia et al., 1992. Abbreviations: m: male, f: female, FWE: family-wise error rate, NKI RS: Nathan Kline Institute Rockland Sample, HCP: Human connectome project.

| Symptom of interest | Anosognosia for hemiplegia | Thalamic aphasia | Post-stroke epilepsy |
| --- | --- | --- | --- |
| Number of patients: total (with/without symptom) | 49 (27/22) | 101 (17/84) | 200 (45/155) |
| Etiology | Ischemic stroke | Ischemic stroke | Ischemic and hemorrhagic stroke |
| Behavioral test | Bisiach anosognosia scale > 1 | Chart review and Aphasia Check List | Chart review or screening questionnaire for detection of epileptic seizures<sup>22</sup> |
| Sex: m/f | 28/21 | 61/40 | 111/90 |
| Age | 30 – 83 (range) | 64.1 ± 14.6 (mean ± sd) | 67.6 ± 13.1 (mean ± sd) |
| Normative cohort | NKI RS, n=50 | NKI RS, n=65 | HCP, n=98 |
| Statistical test for LNM | Two-sample t-test | Two-sample t-test | Two-sample t-test |
| Inference | Random permutation test (voxel level) | Random permutation test (cluster level) | Random permutation test (voxel level) |
| Alpha | $p_{\text{FWE}}(\text{voxel}) < 0.05$ | $p(\text{voxel}) < 0.001$; $p_{\text{FWE}}(\text{cluster}) < 0.05$ | $p_{\text{FWE}}(\text{voxel}) < 0.05$ |
| Main Finding | Right Hippocampus | Left hemisphere language network | Bilateral sensorimotor network |

**Figure 1.**
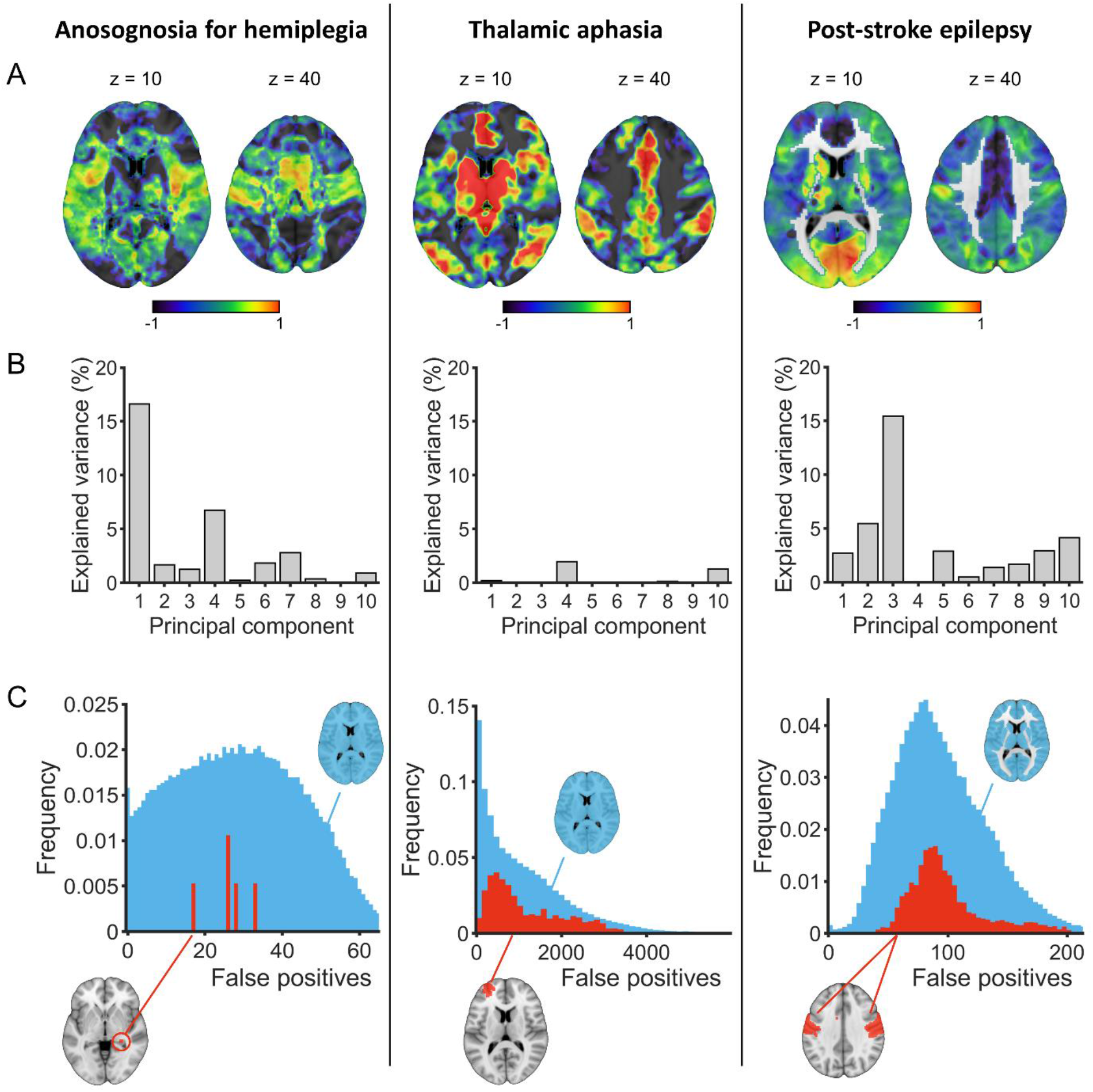
Spatial bias in lesion network mapping. Panel A displays spatial bias operationalized as the pattern of false positives based on 4,000,000 random permutations. The voxelwise frequency of false positives is displayed in a logarithmic fashion relative to the expected value, i.e. log10(fp/mean(fp)). Values > 0 (warm colors) indicate a false positive rate above chance and vice versa for values < 0 (cool colors). Panel B displays the variance of the spatial bias explained by the first 10 principal components of the underlying functional connectome. Panel C displays the distribution of false positives per voxel across all permutations. The distributions are shown for the entire brain mask (light blue) and for voxels significantly associated with the respective symptom (red; arbitrarily scaled for visualization). The sparse histogram for the anosognosia cohort is caused by the significant result only encompassing five voxels. Localizations of significant voxels are displayed on a single representative axial slice (in red as well). Left hemisphere is displayed left.

### Spatial bias estimation and attribution

All analyses were conducted using MATLAB 9.5.0.944444 (R2018b), SPM12 (version 7771), afxRs (version 3.4, https://github.com/afx1337/afxRsPub) and afxStat (version 1.3, https://github.com/afx1337/afxStat) which is based on NiiStat (https://github.com/neurolabusc/NiiStat).

Spatial bias was estimated using a two-step permutation approach. First, 5,000 random permutations of group labels (symptom +/−) were performed within the general linear model to determine a significance threshold. Based on this threshold, an additional 4,000,000 (anosognosia cohort: 6,000,000) permutations were conducted to sample the spatial distribution of false positives under the null hypothesis. This extensive sampling enabled characterization of LNM’s spatial bias independently of any symptom-specific effects.

Similarity of the spatial bias maps across the three independent LNM studies was quantified using linear correlations to assess the extent to which their spatial distributions were consistent. Such consistency would be expected if the observed patterns of LNM reflected fundamental properties of the functional connectome.

To further evaluate whether the observed spatial bias could be attributed to intrinsic features of the connectome, the spatial distribution of false positives was compared with the dominant patterns of the underlying functional connectome. The connectome was represented by connectivity profiles of 1,000 atlas-defined regions.^23^ Dominant patterns were captured using principal component analysis (PCA) of these connectivity maps, and the spatial loadings of the first 10 principal components (PCs) were used to determine how much of the variance in the false-positive maps could be explained by the connectome. This analysis thus quantifies the extent to which the observed spatial bias can be attributed to dominant connectome organization.

Finally, a descriptive analysis localized the actual symptom-associated results within the distribution of false-positive rates (i.e. spatial bias). Estimated false-positive rates, both across all voxels and restricted to significant voxels, were visualized using histograms.

To evaluate the stability of the results with respect to the number of permutations, the analysis was repeated in a split-half approach, computing spatial bias maps separately for the first and last 2,000,000 (anosognosia cohort: 3,000,000) permutations.

All calculations were constrained to a whole brain mask for the anosognosia and thalamic aphasia cohort. Calculations for the post-stroke epilepsy cohort were constrained to a gray matter mask to reduce the high computational demand (caused by the larger number of patients) in the permutative false positive sampling.

## Results

Spatial bias maps, reflected by the distribution of false positives across permutations, are shown in Figure 1A and SI Figure 2. In the anosognosia cohort, spatial bias was found most prominently in the territory of the middle cerebral artery. In the thalamic aphasia cohort, spatial bias was present especially in subcortical structures as well as in several cortical regions. In the post-stroke epilepsy cohort, spatial bias was most pronounced in medial occipital cortex. Overall, these maps exhibited no similarity across the three independent LNM studies, with R^2^ values ranging from 0.4%–0.8% (Table 3), indicating that the spatial distribution of false positives is independent between datasets. Notably, the map derived from the thalamic aphasia cohort showed a more pronounced bias, with several regions displaying false-positive rates above or below an order of magnitude in relation to the expected value (corresponding to absolute values >1 in Figure 1A).

**Table 3.** Shared variance across spatial bias maps.

|  | **Thalamic aphasia** | **Post-stroke epilepsy** |
| --- | --- | --- |
| **Anosognosia for hemiplegia** | 0.8 % | 0.7 % |
| **Thalamic aphasia** | - | 0.4 % |

To characterize dominant spatial patterns in the underlying connectomes, principal component analysis was performed on 1,000 atlas-based connectivity maps. Figures of the spatial loadings for the first 10 components (SI Figure 3–5) and cumulative explained variance plots (SI Figure 6) are provided in the Supplementary Information. The first 10 PCs captured the dominant connectivity patterns, respectively accounting for 88% (anosognosia), 85% (thalamic aphasia), and 91% (post-stroke epilepsy) of the variance. To quantify the extent to which the spatial bias of false positives could be attributed to dominant connectome patterns, these first 10 PCs were used to model the false-positive maps. Across the three cohorts, these components cumulatively accounted for 32% (anosognosia), 4% (thalamic aphasia), and 37% (post-stroke epilepsy) of the variance in the spatial bias, respectively. However, no consistent relation between order of PCs and amount of explained variance was observed (Figure 1B).

To localize the actual symptom-associated findings within the distribution of false positives, estimated false-positive rates were visualized using histograms. Separate histograms were generated for all voxels and for voxels reaching significance (Figure 1C). These histograms showed that the distribution within symptom-associated significant voxels closely mirrored the overall brain-wide distribution. Most importantly, significant voxels were not predominantly located in regions characterized by elevated false-positive rates (i.e. high-bias regions).

Split-half analyses, performed separately on the spatial bias maps obtained from the first and last half of the permutations, produced stable results with only marginal deviations for all analyses presented above. These results indicate that the observed relationships are stable with respect to the number of permutations.

## Discussion

In this study, we investigated the spatial bias inherent to LNM and its relation to dominant features of the underlying connectome across multiple real-world datasets. All analyses were based on group comparisons and rigorous inferential statistics. The spatial bias maps derived from 4,000,000 random permutations showed spatially uneven patterns of false positives reflecting some amount of spatial bias in each individual of the three studied cohorts. However, minimal correspondence of these patterns of spatial bias across the three cohorts, with shared variance as low as 0.4%–0.8% was found. This provides strong evidence that the spatial distribution of false positives is cohort-specific and not systematically determined by the organization of the normative functional connectome. Thus, sources of spatial bias are likely cohort-specific factors, in particular the distribution of lesions within each cohort. The thalamic aphasia cohort showed pronounced levels of false positives, likely reflecting the use of cluster-based statistical inference in conjunction with relatively homogenous lesion locations. In addition, other methodological factors—such as non-stationarity in the data—may also contribute to the observed spatial bias.^24^

Principal component decomposition of the underlying connectome data revealed that dominant connectome features accounted for 4%–37% of the observed spatial bias. Notably, the principal components contributing most strongly to the spatial bias were not those explaining the largest proportion of variance in the connectome, arguing against a systematic relationship between connectome structure and spatial bias patterns.

Finally, distribution analyses using histograms localized the symptom-associated findings within the context of the null distribution of false positives (i.e. spatial bias). Across all three cohorts, the significant results were tightly centered within the overall distribution, with no indication that they were systematically located in regions characterized by high spatial bias. This observation suggests that the variance explained by behavioral variables predominates over the inherent structures captured by the data under the null model, highlighting that symptom-related effects emerge distinctly from the cohort-specific spatial bias inherent to LNM.

Further contextualization of the present results is challenging. The only other study known to us that examined spatial distributions of false positives—albeit using a different methodological approach— found a pronounced spatial bias in functional fMRI data, particularly in the posterior cingulate cortex. This effect was attributed to non-stationarity and locally increased smoothness in the data.^25^ In our post-stroke epilepsy cohort, false positives were found most pronounced in a nearby region encompassing cuneus and calcarine sulcus. However, no meaningful relationship between spatial bias and local smoothness (computed from the corresponding LNMs)^26^ was found in an additional exploratory analysis (R^2^ < 0.1 %).

These findings together highlight that spatial bias is a measurable feature in LNM, but one that is cohort-specific and independent of the dominant patterns of the normative connectome and not pivotal for symptom-associated LNM findings.

The previous study by van den Heuvel et al. provided the initial impetus for the present work through its hypothesis of systematic spatial bias in lesion network mapping.^9^ Both studies share the approach of examining spatial bias across multiple cohorts. However, a key distinction is that, unlike van den Heuvel et al., who compared unthresholded patterns directly, the current study is the first to apply rigorous permutation-based inference to sample the spatial bias. This thresholded, statistically controlled approach allows for a rigorous evaluation of false-positive distributions and their relationship to underlying connectivity patterns. Additionally, this approach more closely reflects the recommended methodological standards for neuroimaging studies^27^ and therefore allows spatial bias to be evaluated under conditions that approximate real-world LNM analyses.^11–17^ Taken together, the evidence from both studies suggests that the validity of lesion network mapping critically depends on methodological factors, particularly the level and rigor of statistical analysis. In our view, robust results are most likely to emerge from study designs that include an explicit control group or an implicit model-based control (e.g., regression with behavioral data), combined with the application of appropriate inferential frameworks such as permutation-based tests.

Although the present study provides strong evidence against the existence of a methodologically driven spatial bias related to connectome features, several limitations should be acknowledged. These include the limited number of cohorts analyzed, the use of different normative connectomes, and variation in inferential approaches (voxel-versus cluster-level analyses). Nevertheless, given the numerically extremely low similarity of the spatial bias maps across cohorts (0.4–0.8%), it seems unlikely that the inclusion of additional cohorts would fundamentally alter the overall conclusion.

## Conclusion

In conclusion, our results indicate that spatial bias in lesion network mapping is largely cohort-specific and not systematically determined by dominant features of the normative connectome. Rigorous study designs combined with appropriate inferential frameworks yield robust, interpretable results and minimize the risk of bias. Our findings support the validity of LNM for investigating disorder-related networks while highlighting the critical importance of methodological rigor in study design and statistical analysis.

## Non-Standard Abbreviations

LNM: lesion network map(ping)
PCA: Principal component analysis
PC(s): Principal component(s)

## Author contributions

Conceptualization: MW, TR, AS; Software, Methodology, Formal Analysis, Writing – Original Draft: MW; Writing – Review & Editing: all authors. Large language models were used solely for language editing.

## References

1. Avena-Koenigsberger A, Misic B, Sporns O. Communication dynamics in complex brain networks. Nature reviews. Neuroscience 2017;19(1):17–33.

2. Michalopoulou PG, Meshreky KM, Hommerich Z, Shergill SS. Neuromodulation and neural networks in psychiatric disorders: current status and emerging prospects. Psychological medicine 2025;55:e281.

3. Sutterer MJ, Tranel D. Neuropsychology and cognitive neuroscience in the fMRI era: A recapitulation of localizationist and connectionist views. Neuropsychology 2017;31(8):972–980.

4. Boes AD, Prasad S, Liu H, Liu Q, Pascual-Leone A, Caviness VS, et al. Network localization of neurological symptoms from focal brain lesions. Brain : a journal of neurology 2015;138(Pt 10):3061–3075.

5. Fox MD. Mapping Symptoms to Brain Networks with the Human Connectome. The New England journal of medicine 2018;379(23):2237–2245.

6. Joutsa J, Corp DT, Fox MD. Lesion network mapping for symptom localization: recent developments and future directions. Current opinion in neurology 2022;35(4):453–459.

7. Sperber C, Dadashi A. The influence of sample size and arbitrary statistical thresholds in lesion-network mapping. Brain : a journal of neurology 2020;143(5):e40.

8. Cohen AL, Fox MD. Reply: The influence of sample size and arbitrary statistical thresholds in lesion-network mapping. Brain : a journal of neurology 2020;143(5):e41.

9. van den Heuvel MP, Libedinsky I, Quiroz Monnens S, Repple J, Sommer I, Cocchi L. Investigating the methodological foundation of lesion network mapping. Nature neuroscience 2026.

10. Siddiqi SH, Horn A, Schaper FL, Khosravani S, Cohen AL, Joutsa J, et al. The methodological foundations of lesion network mapping remain sound. bioRxiv 2026.

11. Wawrzyniak M, Klingbeil J, Zeller D, Saur D, Classen J. The neuronal network involved in self-attribution of an artificial hand: A lesion network-symptom-mapping study. NeuroImage 2018;166:317–324.

12. Klingbeil J, Wawrzyniak M, Stockert A, Karnath H-O, Saur D. Hippocampal diaschisis contributes to anosognosia for hemiplegia: Evidence from lesion network-symptom-mapping. NeuroImage 2020;208:116485.

13. Stockert A, Hormig-Rauber S, Wawrzyniak M, Klingbeil J, Schneider HR, Pirlich M, et al. Involvement of Thalamocortical Networks in Patients With Poststroke Thalamic Aphasia. Neurology 2023;100(5):e485–e496.

14. Schaper FLWVJ, Nordberg J, Cohen AL, Lin C, Hsu J, Horn A, et al. Mapping Lesion-Related Epilepsy to a Human Brain Network. JAMA neurology 2023;80(9):891–902.

15. Al-Fatly B, Neudorfer C, Kaski D, Lang AE, Kühn AA, Fox MD, et al. Lesion network of oculogyric crises maps to brain dopaminergic transcriptomic signature. Brain : a journal of neurology 2024;147(6):1975–1981.

16. Klingbeil J, Wawrzyniak M, Stockert A, Brandt M-L, Schneider H-R, Metelmann M, et al. Pathological laughter and crying: insights from lesion network-symptom-mapping. Brain : a journal of neurology 2021;144(10):3264–3276.

17. Rangus I, Rios AS, Horn A, Fritsch M, Khalil A, Villringer K, et al. Fronto-thalamic networks and the left ventral thalamic nuclei play a key role in aphasia after thalamic stroke. Communications biology 2024;7(1):700.

18. Sutterer MJ, Bruss J, Boes AD, Voss MW, Bechara A, Tranel D. Canceled connections: Lesion-derived network mapping helps explain differences in performance on a complex decision-making task. Cortex; a journal devoted to the study of the nervous system and behavior 2016;78:31–43.

19. Siddiqi SH, Philip NS, Palm ST, Carreon DM, Arulpragasam AR, Barredo J, et al. A potential target for noninvasive neuromodulation of PTSD symptoms derived from focal brain lesions in veterans. Nature neuroscience 2024;27(11):2231–2239.

20. Karnath H-O, Sperber C, Rorden C. Mapping human brain lesions and their functional consequences. NeuroImage 2018;165:180–189.

21. Nichols TE, Holmes AP. Nonparametric permutation tests for functional neuroimaging: a primer with examples. Human brain mapping 2002;15(1):1–25.

22. Placencia M, Sander JW, Shorvon SD, Ellison RH, Cascante SM. Validation of a screening questionnaire for the detection of epileptic seizures in epidemiological studies. Brain : a journal of neurology 1992;115 (Pt 3):783–794.

23. Schaefer A, Kong R, Gordon EM, Laumann TO, Zuo X-N, Holmes AJ, et al. Local-Global Parcellation of the Human Cerebral Cortex from Intrinsic Functional Connectivity MRI. Cerebral cortex (New York, N.Y. : 1991) 2018;28(9):3095–3114.

24. Hayasaka S, Phan KL, Liberzon I, Worsley KJ, Nichols TE. Nonstationary cluster-size inference with random field and permutation methods. NeuroImage 2004;22(2):676–687.

25. Eklund A, Nichols TE, Knutsson H. Cluster failure: Why fMRI inferences for spatial extent have inflated false-positive rates. Proceedings of the National Academy of Sciences of the United States of America 2016;113(28):7900–7905.

26. Worsley KJ, Marrett S, Neelin P, Vandal AC, Friston KJ, Evans AC. A unified statistical approach for determining significant signals in images of cerebral activation. Human brain mapping 1996;4(1):58–73.

27. Roiser JP, Linden DE, Gorno-Tempinin ML, Moran RJ, Dickerson BC, Grafton ST. Minimum statistical standards for submissions to Neuroimage: Clinical. NeuroImage. Clinical 2016;12:1045–1047.

