## Supplementary Information for "Spatial Bias in Lesion Network Mapping Is Connectome-Independent"

### Supplementary Figures

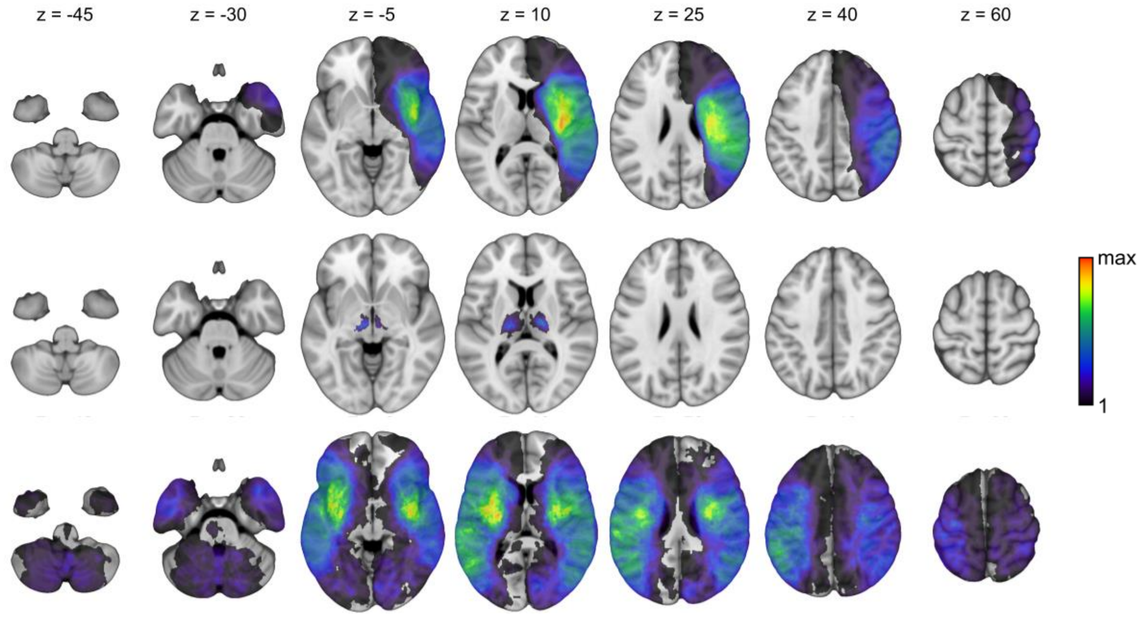


**SI Figure 1. Lesion overlays.** Lesion overlays for anosognosia for hemiplegia cohort (first row), thalamic aphasia cohort (middle row) and post-stroke epilepsy cohort (last row). Coordinates refer to MNI space in mm. Left hemisphere is displayed left.

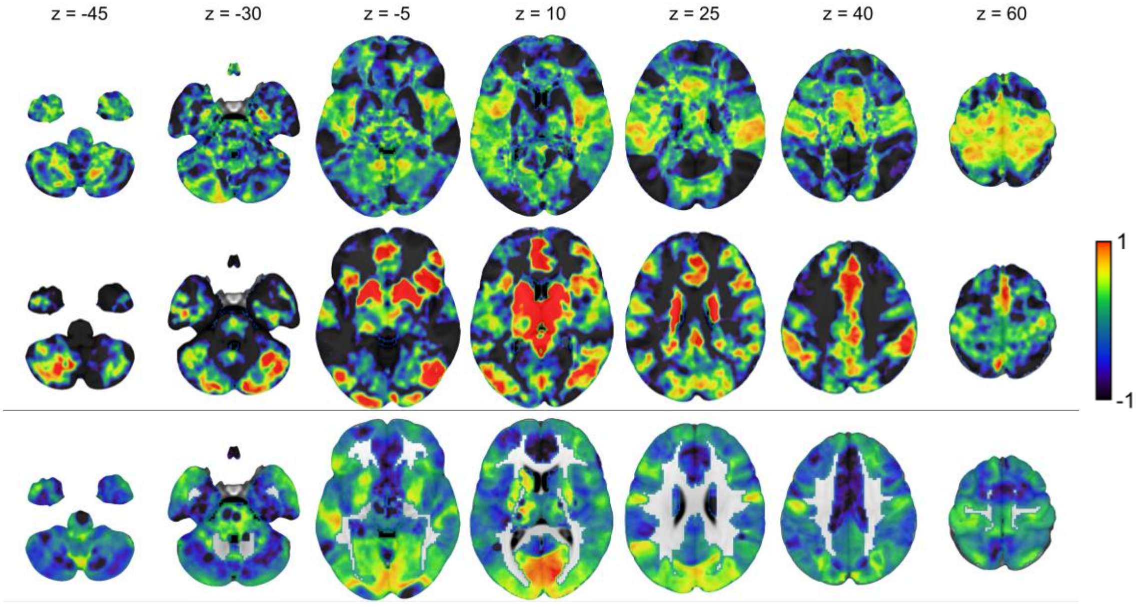


**SI Figure 2. Spatial bias in lesion network mapping.** Spatial bias is displayed for the cohorts examining anosognosia for hemiplegia (first row), thalamic aphasia (middle row) and post-stroke epilepsy (last row). Spatial bias is operationalized as the pattern of false positives based on 4,000,000 random permutations. The voxelwise frequency of false positives is displayed in a logarithmic fashion relative to the expected value, i.e. log10(fp/mean(fp)). This way, values > 0 indicate a false positive rate above chance and vice versa for values < 0. Coordinates refer to MNI space in mm. Left hemisphere is displayed left.

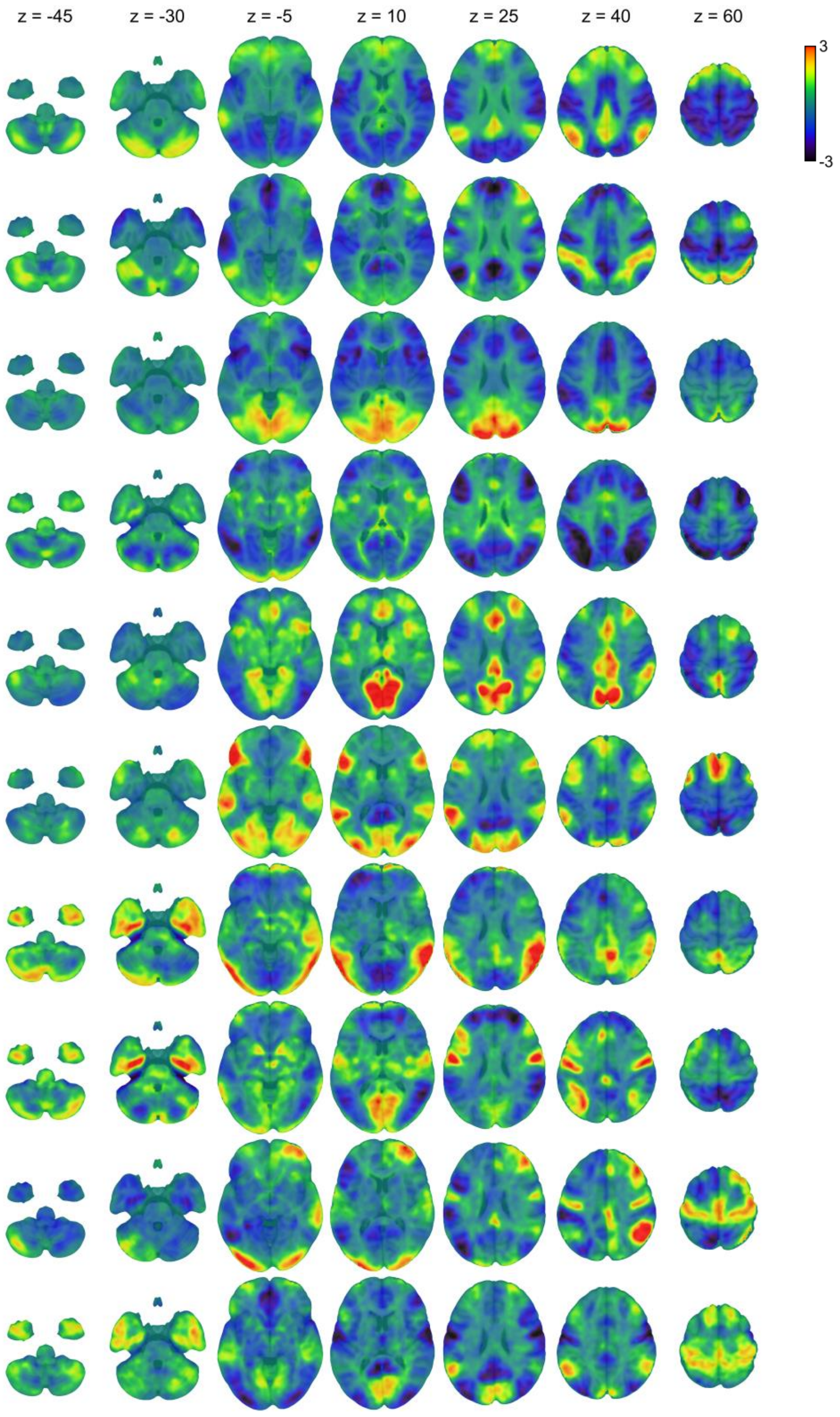


**SI Figure 3. Principal components of the functional connectome (anosognosia cohort).** The normative functional connectome, represented by connectivity profiles of 1,000 atlas-defined seed regions, was decomposed using principal component analysis. Each row sows the spatial loadings of the respective principal component, scaled to unit variance and rendered on an MNI template. Eigenvector signs were fixed to resolve sign indeterminacy by requiring the element with the largest absolute magnitude to be positive, and the spatial loadings inherit this orientation. Left hemisphere is displayed on the left.

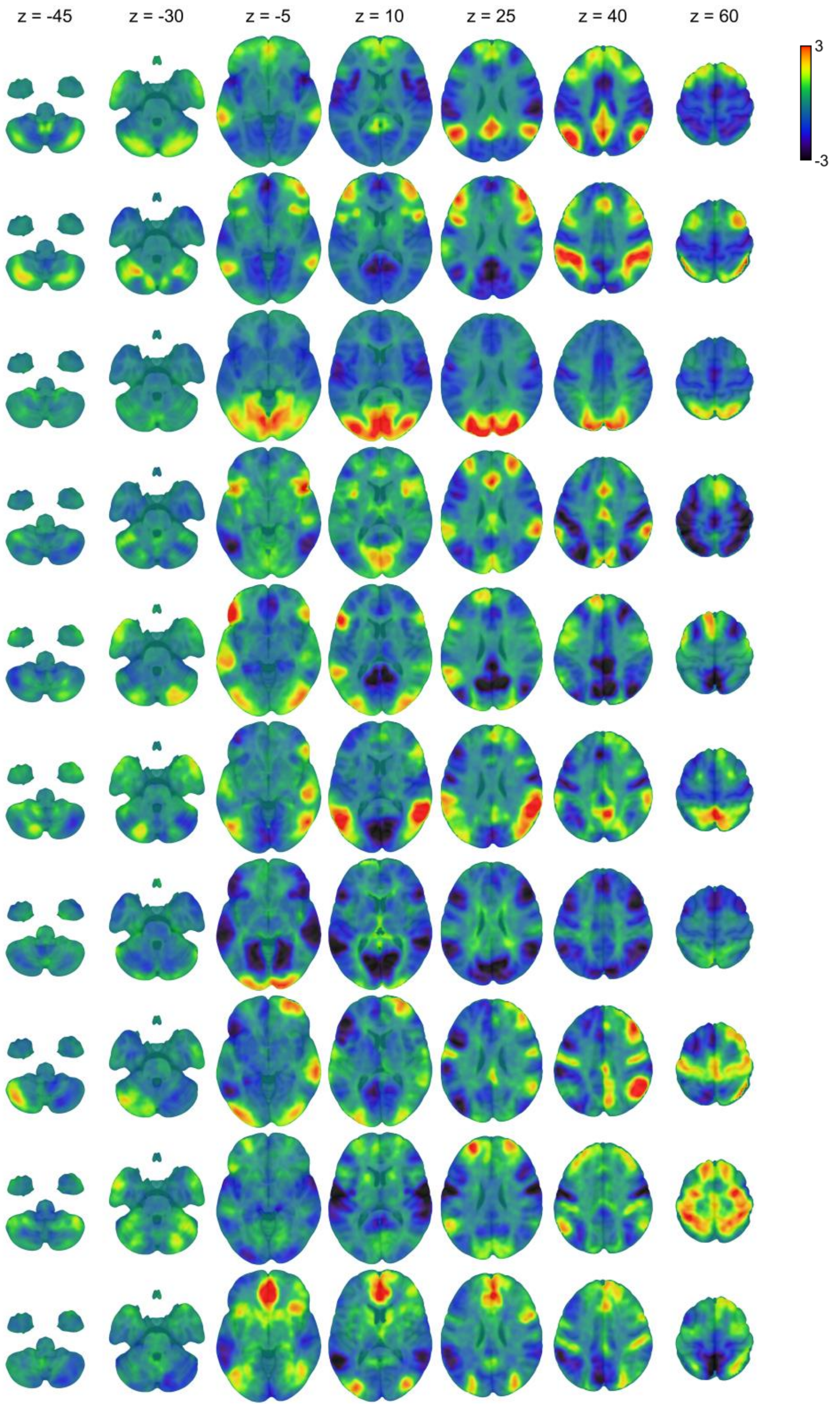


**SI Figure 4. Principal components of the functional connectome (thalamic aphasia cohort).** The normative functional connectome, represented by connectivity profiles of 1,000 atlas-defined seed regions, was decomposed using principal component analysis. Each row sows the spatial loadings of the respective principal component, scaled to unit variance and rendered on an MNI template. Eigenvector signs were fixed to resolve sign indeterminacy by requiring the element with the largest absolute magnitude to be positive, and the spatial loadings inherit this orientation. Left hemisphere is displayed on the left.

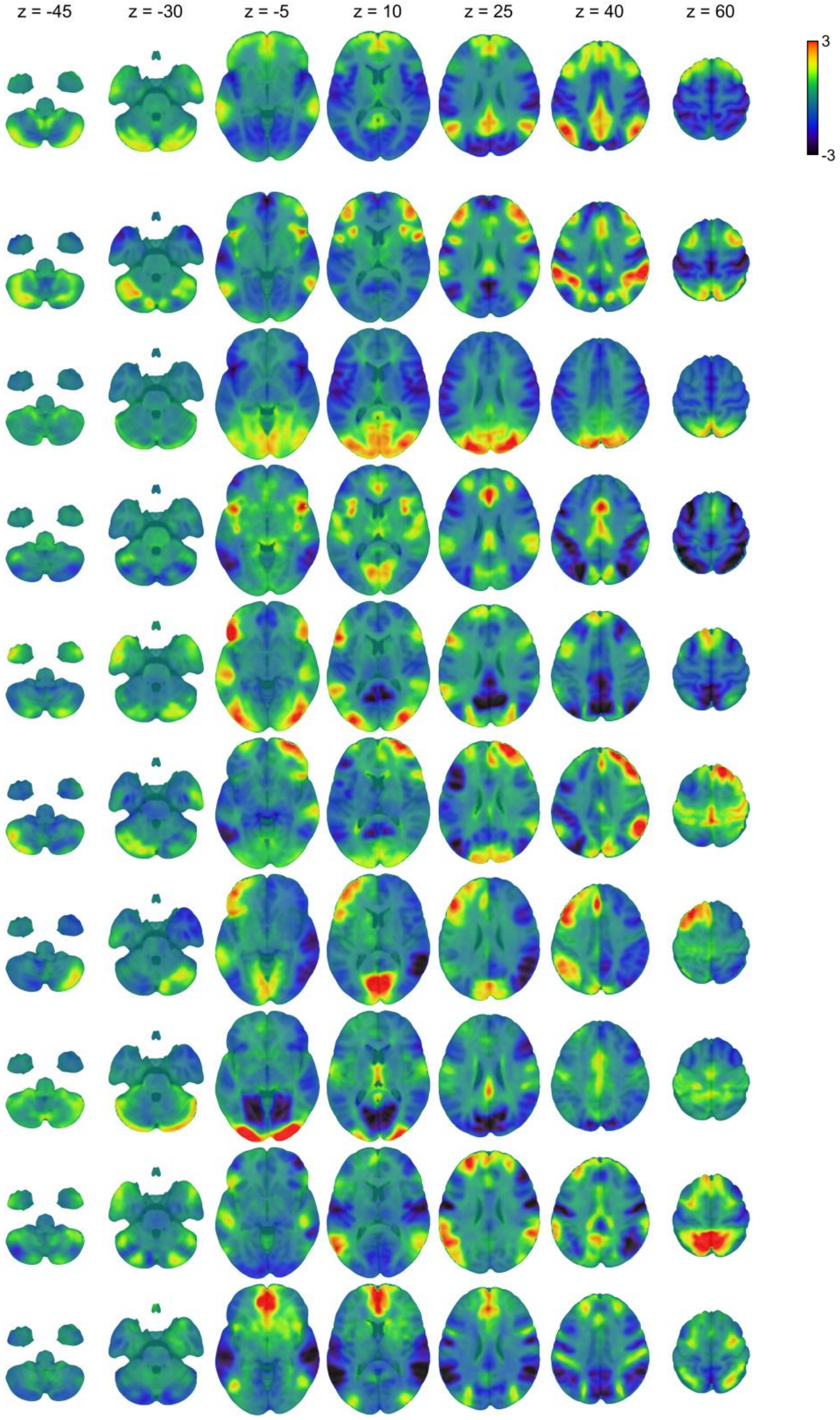


**SI Figure 5. Principal components of the functional connectome (post-stroke epilepsy cohort).** The normative functional connectome, represented by connectivity profiles of 1,000 atlas-defined seed regions, was decomposed using principal component analysis. Each row sows the spatial loadings of the respective principal component, scaled to unit variance and rendered on an MNI template. Eigenvector signs were fixed to resolve sign indeterminacy by requiring the element with the largest absolute magnitude to be positive, and the spatial loadings inherit this orientation. Left hemisphere is displayed on the left.

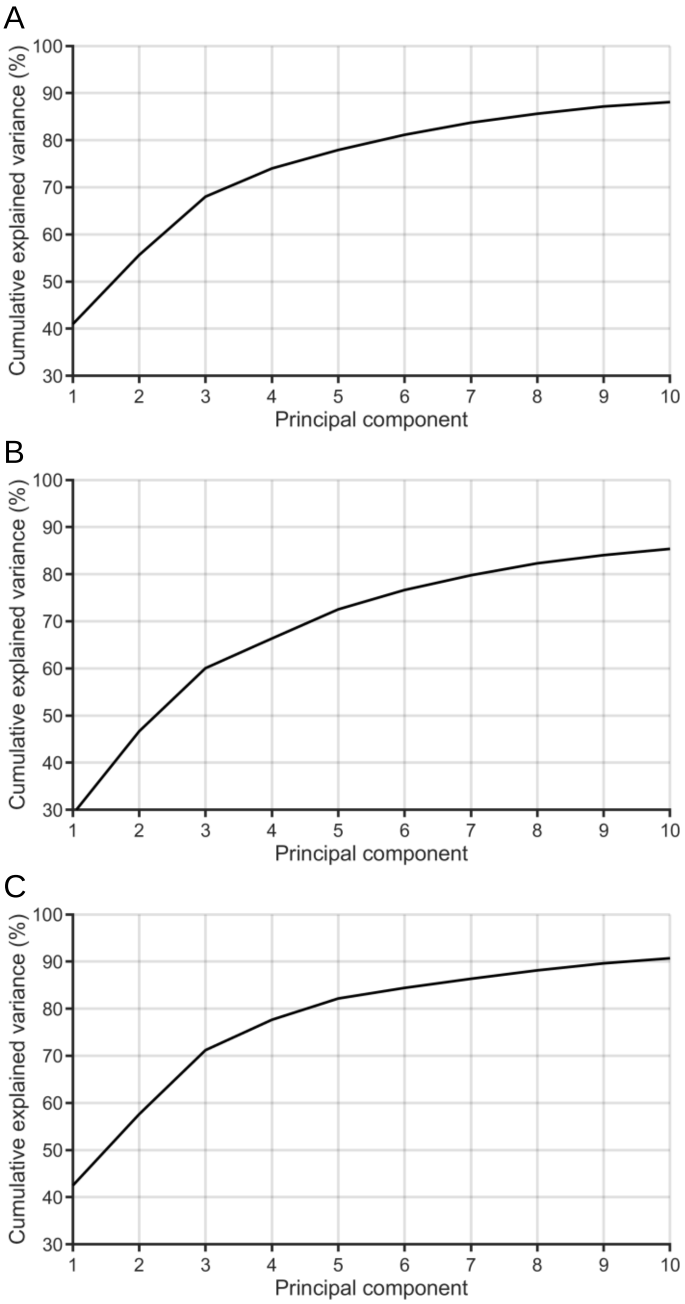


**SI Figure 6. Cumulative explained variance.** The plots show the cumulative explained variance of the first 10 principal components of the functional connectome, represented by connectivity profiles of 1,000 atlas-defined seed regions. Results are shown for the normative connectomes used in the analyses of (A) anosognosia for hemiplegia, (B) thalamic aphasia, and (C) post-stroke epilepsy.
